# Polysomal profiling coupled to allele-specific proteomics reveals an EIF4H tranSNP allele possessing higher mRNA translation potential

**DOI:** 10.1101/2025.04.17.649328

**Authors:** Meriem Hadjer Hamadou, Laura Alunno, Daniele Peroni, Michael Pancher, Tecla Venturelli, Romina Belli, Erik Dassi, Alessandro Romanel, Alberto Inga

**Author notes:** equal last and corresponding authors. equal first authors (Meriem Hadjer Hamadou, Laura Alunno, Daniele Peroni).

## Abstract

To search for genetic sources of allele-specific mRNA translation, we leveraged heterozygous polymorphisms and variants present in the exome of HCT116-derived cell lines, computing allelic fractions in total and polysome-associated RNA from RNA-seq data. Allelic imbalance in polysomal RNA led us to nominate 52 coding variants associated with allele-specific mRNA translation, of which 16 are nonsynonymous. To validate instances of allele-specific translation, a proof-of-concept proteomics approach was developed to quantify the relative expression of pairs of endogenous proteins resulting from the decoding of alleles containing the nonsynonymous heterozygous variants. In particular, the G>A, R183H missense SNV rs1554710467 in the EIF4H gene was investigated. The alternative peptide containing H183 was significantly more abundant than the corresponding one containing R183, consistent with the overrepresentation of the alternative allele in polysomal RNA in HCT116 cells. A dual-fluorescence ribosome-stalling assay confirmed the enhanced translation potential of the variant allele. The two EIF4H allelic proteins exhibited similar stability and subcellular localization. These findings support the classification of rs1554710467 as a gain-of-function allele. This study demonstrates the feasibility of using allele-specific proteomics at the endogenous protein levels, exploiting heterozygous coding variants. Overall, our approach extends the toolbox available to investigate allele-specific differences in mRNA translation potential, a relatively underexplored layer of gene expression control that could underlie inter-individual differences in disease-relevant phenotypes.

## Introduction

Translational control of mRNAs has been consistently revealed as a crucial process in cancer (Silvera et al. 2010; Robichaud et al. 2019), yet there are relatively few studies focusing on the identification of germline and somatic sources of inter-individual variation in the fate of specific cancer-relevant transcripts. To bridge this gap, we recently developed a pipeline tune to discover alleles associated with differences in mRNA translation potential (Valentini et al. 2021; Hamadou et al. 2025). Our approach relies on polysome isolation via sucrose-gradient fractionation followed by RNA extraction and sequencing (Chassé et al. 2017; Zaccara et al. 2014; Zuccotti and Modelska 2016; Wang et al. 2020; King and Gerber 2016). Heterozygous SNPs and SNVs that are present in mRNAs are evaluated by computing allelic fractions (AF) for polysome-bound RNA samples that are compared to the AF measured from total RNA and by annotating variations (delta AF) that are consistent among biological replicates and/or above experimental variance (Valentini et al. 2021; Hamadou et al. 2025). The method relies on the assumption that the translatome, *i.e.* the repertoire of mRNAs bound to polysomes, is a proxy of the proteome so that there is a direct proportionality between the translation efficiency of a given allele and its relative association with polysomes. In our previous studies, we focused on validating candidate SNPs, labeled as tranSNPs, that are located in the untranslated regions of mRNAs and for which we could also generate predictions of trans factors binding, namely RNA-Binding Proteins (RBPs). Candidates were prioritized based on the magnitude and consistency of variations in AF, the frequency of the minor alleles in human populations, and the relevance of the gene for cancer, with a particular interest in p53-dependent responses. Validation experiments comprised orthogonal approaches to calculate AF, cloning of UTR sequences in reporter plasmids to assess functional differences among the two alleles, and genome editing to derive potentially isogenic trios of genotypes for functional analysis at the endogenous gene level (Valentini et al. 2021; Hamadou et al. 2025). In the case of the rs1053639 T/A tranSNP in the 3’UTR of the DDIT4 gene, an inhibitor of mTORC1 (Coronel et al. 2022; Whitney et al. 2009), the T allele was over-represented in the polysomes in heterozygous cells, and the derived T-homozygous clones exhibited higher DDIT4 protein levels and showed more effective repression of mTORC1 responses upon ER stress. A-homozygous clones instead proliferated more in competition assays or when injected in zebrafish embryos.

In this study, we focused on tranSNPs resulting in missense changes and developed a proof-of-concept approach to validate them by proteomics. Briefly, we compared whole proteome, fractionation, and targeted proteomics with or without the use of spiked-in heavy-isotope references to explore the possibility of quantifying bi-allelic peptides resulting from heterozygous nonsynonymous polymorphisms and variants.

Although the impact of genetic factors on the human proteome is still largely unexplored, previous proteomics studies have explored the possibility of identifying genetic variants associated with circulating protein concentrations, protein quantitative trait loci (pQTL, cis-pQTL), cancer-associated mutations, and developed methods to quantify proteomes across various biological samples (Yao et al. 2018; He et al. 2020; Johansson et al. 2013; Wu et al. 2013; Shi et al. 2018; Tan et al. 2017; Ogata et al. 2022; Lin et al. 2023; Tan et al. 2020; Wang et al. 2011).

Our approach differs as it exploits biallelic genetic variants present in the exome of a model cell line and uses a filtering approach based on allelic fractions in polysome-bound RNAs from RNA-seq to identify and prioritize candidate genetic sources (tranSNPs) of variation in protein levels. In our proof-of-concept study, peptide-encompassing tranSNP positions could be investigated for three targets, and in all cases, imbalances between reference or alternative peptides were observed, and these were consistent with the delta-AF measured in the polysomal RNA-seq. In particular, we validated the rs1554710467 SNV present in HCT116 cells that results in the heterozygous expression of the R183H mutant of EIF4H. Results established that the variant mRNA allele that is over-represented in polysomal RNA fractions is associated with a higher abundance of the corresponding peptide (single amino acid variant, H183 vs R183). The difference is not dependent on a change in protein half-life and is not associated with altered subcellular localization of EIF4H, suggesting that the SNV represents a gain-of-function allele resulting in higher EIF4H protein levels.

## Materials and Methods

### Polysome profiling, RNA extraction, and Sanger sequencing

HCT116 cells grown in 150 mm dishes till 80% confluence were incubated with 50 µg /ml cycloheximide at 37°C for 10 min to immobilize the ribosomes on mRNAs Cells were rinsed with ice-cold PBS containing 50 µg /ml cycloheximide and then gently scraped using about 600 µl of ice-cold lysis buffer (20 mM Tris-HCl pH 7.5, 100 mM KCl, 5 mM MgCl_2_, 0.5 % Nonidet P-40, 0.2 U/µl RNasin Ribonuclease Inhibitor [Promega], cOmplete, Mini Protease Inhibitor Cocktail 1X [Roche], 100 µg /ml cycloheximide). The lysates were then incubated on ice for 10 min and centrifuged for 10 min at 12,000 *x g* at 4 °C to remove the nuclei. The cytoplasmic lysates were loaded onto a 15-50% linear sucrose gradient prepared in salt buffer (100 mM NaCl, 20 mM Tris-HCl pH7.5, and 5 mM MgCl2) and separated to equilibrium density by ultracentrifugation using a Beckman SW41 rotor at 40.000 rpm for 1 hour and 40 minutes at 4°C. The sucrose gradients were then fractionated using a Teledyne Isco model 160 gradient analyzer equipped with a UA-6 UV/VIS detector, monitoring the absorbance at 254 nm. Thirteen fractions of 1 ml each were isolated using a 50% sucrose solution pumped at the bottom of the centrifuged tube. RNA was harvested from pooled fractions corresponding to either subpolysomal subunits (SUB), 80S monosome, light or heavy polysomes (POL) using the TRIzol reagent (Invitrogen ^TM^). An aliquot of the unfractionated cytoplasmic lysate was processed to obtain total RNA (TOT). The RNA was retro-transcribed, and the sequences containing the candidate tranSNPs were amplified by PCR using AmpliTaq Gold 360 Master Mix (Thermo Fisher Scientific) and specific primers (**Table S4**). The PCR products were purified by the spinNAker GEL&PCR DNA purification kit (Euroclone) and subjected to Sanger sequencing by Mix2Seq Kit (Eurofins Genomics). Analysis of the electropherograms was performed by Fiji. software. Quantification was performed by assessing the area under the peaks of the tranSNP nucleotides as a metric to calculate allelic fraction. The normalization process considered the relative efficiency of the four nucleotides’ incorporation in the sequence context surrounding the SNP position. The allelic fraction for targeted genes was determined for at least three independent experiments. Student’s t-test was used to calculate the p-value.

### Western blot and cycloheximide-chase assay

Cytoplasmic lysates were prepared and centrifuged on sucrose gradients and fractionated as described above. Proteins were isolated from the same volume from each fraction using a RIPA buffer containing protease inhibitor cocktail (cOmplete, Mini Protease Inhibitor Cocktail, Roche). Before western blotting, the lysates were denatured by boiling at 95°C for 10 min in 1x Laemmli Sample Buffer (Bio-Rad), then subjected to SDS-PAGE and transferred to a nitrocellulose membrane (Cytiva Amersham™ Reinforced Nitrocellulose Blotting Membrane). Each membrane was blocked with 5% dry milk in phosphate-buffer saline and 0.1% Tween-20 (PBST) for 1 h and then incubated with the indicated primary antibodies overnight at 4°C. The membranes were rinsed and incubated with peroxidase-conjugated secondary antibodies for 1 hour at RT. Secondary antibodies were used at 1:10000 dilutions. Membranes were developed with ECL^TM^ Select western blot detection reagents (Amersham Biosciences, UK) and imaged on the ChemiDoc XRS imaging system (Bio-Rad). Image processing and densitometric quantification of the bands were performed using the Image Lab software. Antibodies used for western blot analysis are listed in **Table S5**. To estimate EIF4H protein half-life, 6×10^5^ HCT116 cells were seeded in a 6-well plate format. After 24 hours, cells from one well were collected by trypsinization, and the cell pellet was resuspended in 50 µl of RIPA lysis buffer supplemented with 1X protease inhibitors (Roche) to prepare the t0 sample, which was kept at −20C. In the meantime, the media was aspirated from the remaining wells and replaced with fresh media containing 100 μg/ml of cycloheximide (Cayman Chemical) to halt protein synthesis. After 2,4,6, and 8 hours, cells were pelleted from the wells and lysed as the t0 sample. Once all the pellets were harvested, proteins were extracted by centrifugation at 15000 rpm for 20 minutes at 4°C, quantified by the Bicinchoninic Acid Assay (Thermo Scientific Pierce BCA Protein Assay Kit), and a Western Blot was performed. 35 µg of time point t0 protein extract was used for the Western Blot. For the other timepoints, the protein amounts were loaded volumetrically to the reference t0. EIF4H and β-actin antibodies were used as primary antibodies. β-actin was found to be a stable protein in the time course of the experiment and thus used as a loading control even in this context.

### Sample Preparation for proteomic analysis

For whole proteome analysis, HCT-116 cells were harvested and lysed in RIPA buffer at 4°C for 30 min, followed by centrifugation to clarify the samples. The protein concentration was determined using a BCA assay. Proteins (100μg) were then reduced by 10 mM dithiothreitol (DTT) at 56*°*C for 30 min and alkylated by 22.5 mM iodoacetamide (IAA) at 25 °C for 30 min in dark. The addition of 10 mM DTT quenched excess IAA. Protein digestion was performed using carboxylate magnetic beads (GE Healthcare) as described before (Hughes et al. 2019). Briefly, washed SP3 beads were added to protein mixtures with a 1:10 ratio (w/w) protein/beads. Subsequently, acetonitrile (ACN) was added to a final concentration of 70% (v/v), and samples were mixed at room temperature for 18 min. The supernatant was removed with the assistance of a magnetic stand, and the beads were rinsed two times with 70% ethanol and once with ACN. Beads were then resuspended in 45 μL of 50 mM NH4HCO3, 5mM CaCl2, supplemented with trypsin (enzyme: protein ratio of 1:20). After overnight digestion at 37 °C under mixing, the peptide mixture was collected by incubation on a magnetic rack, and the supernatant was subjected to SP2 peptide clean-up (Waas et al. 2019). Peptide mixtures were bound to magnetic carboxylate-modified beads (GE Healthcare) in a ratio of 1:15 by the addition of ACN, such that the final ACN concentration was exactly 95%. Beads were then washed twice with 100% ACN, and the peptides were eluted with 50 μL of 2% ACN in water by mixing at 1000 rpm for 10 min at 25 °C. Samples were centrifuged at 13000 rpm for 10 seconds and placed on the magnetic rack. The supernatant was transferred to a clean tube and acidified with formic acid (final concentration of 0.1%) for LC-MS analysis or further processing. To increase proteome coverage, 50 μg of digested peptides were fractionated using the High pH Reversed*-*Phase Peptide Fractionation Kit (Thermo Fisher Scientific). Peptides were diluted to 300 μL with 0.1% trifluoroacetic acid and fractionated as described in the manufacturer’s manual. The resulting ten fractions were then dried by vacuum centrifugation and reconstituted in 0.1% formic acid for LC-MS analysis. To enrich the proteins of interest (EIF4H, RPA1, RBM17), HCT116 cell lysates or TCA-precipitated proteins from RNP and 40S fractions were resolved by SDS polyacrylamide gels electrophoresis using Bolt 10% Bis-Tris minigel (Thermo Fisher Scientific) and stained with Coomassie (Imperial Protein Stain, Thermo Fisher Scientific). For each sample, the gel bands corresponding to the regions 20-30 kDa and 65-85 kDa were excised and cut into small pieces (∼1 mm^3^). The gel pieces were then destained with 50% acetonitrile (ACN) in 100 mM NH_4_HCO_3_, dehydrated by net ACN, and dried in a SpeedVac. Samples were subjected to reduction and alkylation with 10 mM DTT and 55 mM IAA, respectively. Gel pieces were washed repeatedly in 100 mM NH_4_HCO_3_, followed by 100% ACN. After drying, the excised gel pieces were incubated with trypsin (Thermo Fisher Scientific) at 12.5 ng/μl in 100 mM NH_4_HCO_3_ and placed on ice. After 45 min, the digestion continued at 37°C overnight. The next day, the supernatant was transferred into a clean tube, and the peptides were sequentially extracted from the gels with 30% ACN, 3% trifluoroacetic acid (TFA), and 100% ACN. All the supernatants were combined with the solution retrieved after digestion and dried in a SpeedVac. The peptides were then acidified with 1% TFA to a pH <2.5, desalted on homemade C18 stage tips, and resuspended in 0.1% formic acid in water for LC-MS analysis. Cleaned-up peptides from all samples were stored at −20 °C until measurement.

### Liquid chromatography-tandem mass spectrometry (LC-MS/MS) analysis

For shotgun label-free quantification analysis, digested samples were separated using an Easy-nLC 1200 system (Thermo Fisher Scientific) and loaded onto a reversed-phase column (Acclaim PepMap RSLC C18 column 2 µm particle size, 100Å pore size, id 75 µm), heated at 40°C. A two-component mobile phase system consisting of 0.1% formic acid in water (buffer A) and 0.1% formic acid in acetonitrile (buffer B) was used. Peptides were eluted using the following gradient: 5% to 25% buffer B over 52 minutes, 25% to 40% over 8 minutes, 40% to 98% over 10 minutes, followed by 10 minutes at 98% of buffer B. The flow rate was set to 400 nL/min. Samples were injected into an Orbitrap Fusion Tribrid mass spectrometer (Thermo Fisher Scientific, San Jose, CA, USA), and data were acquired in data-dependent mode (2100 V). The ion transfer tube temperature was set to 275°C. Full scans were performed at a resolution of 120.000 FWHM (at 200 m/z) and an AGC target of 1×10^6^. The precursor mass range was 110-1100 m/z, with the first mass for fragments set at 140 m/z and a maximum injection time of 50 ms. The dynamic exclusion filter was enabled with a duration of 30 sec. Each full scan was followed by MS/MS scans (HCD, collision energy of 30%) over a 3-second cycle time, with a maximum injection time of 54 ms (Orbitrap) and an AGC target of 2.5×10^4^. In the ion prioritization method, selected ion masses and charge states were given priority (**Table S4A**). Fractions were acquired using identical MS1 and MS2 parameters as described above. For PRM LC-MS/MS measurements, peptides containing the biallelic amino acid position of interest were targeted. An *in-silico* digestion with trypsin was performed on the selected proteins to identify peptides. Being constrained by the biallelic position, it was not always possible to avoid amino acid modifications, such as the oxidation of methionine. In addition, in the case of the EIF4H protein, the mutation led to a change in peptide length (10 AA in the ALT version versus 8 AA in the REF). In the REF version, a peptide of similar length (i.e., 10 AA, including one missed cleavage) was not observed. When two charge states met the mass range, we selected the better charge state based on a manual check of the mass spectrum signal response, such as the signal-to-noise ratio, the transitions, the peak shape, and the peak area. Heavy reference peptides (Thermo Fisher Scientific) for EIF4H and RPA1 proteins were spiked into digested samples enriched for EIF4H or RPA1 proteins, as described above. Heavy peptides were maintained at a constant concentration of approximately 100 femtomoles in all samples. One microliter of each sample was loaded onto the LC column for analysis. LC settings were identical to those described above, except for the use of a 30 cm reversed-phase long column (inner diameter 75 µm, 1.7 µm particle size, MSWIL, the Netherlands), heated to 40°C, with a flow rate of 200 nL/min. The isolation list for the heavy and light peptides was provided as a mass list with charge state information (**Table S4B**). tMSn scans were acquired in the Orbitrap at R = 60,000, with an AGC target value set to standard and a maximum injection time of 118 ms. HCD collision energy was set to 30%. Ion source parameters were as follows: spray voltage +2100V, and ion transfer tube temperature 200°C. For all acquisitions, a blank and a quality control sample were run between samples to prevent carryover and monitor instrument performance. QCloud was used to assess longitudinal instrument performance throughout the project (Chiva et al. 2018).

### Data analysis

Extracted ion chromatograms for all transitions were inspected using Xcalibur Qual Browser (Thermo Fisher Scientific). For the shotgun analysis, peptide searches were performed using Proteome Discoverer 3.1 software (Thermo Fisher Scientific) against the *Human* proteome (Uniprot, downloaded June 2023) and a database of common contaminants. Proteins were identified using the MASCOT search engine with a mass tolerance of 10 ppm for precursor and 0.02 Da for the product. Trypsin was specified as the digestion enzyme, allowing up to 5 missed cleavages. Carbamidomethylation of cysteine was set as a static modification, while oxidation (M) and acetylation (protein N-term) were considered variable modifications. The false discovery rate (FDR) was filtered at <0.01 at the PSM, peptide, and protein levels. Potential contaminants were excluded from the results. Peak intensities were log2-transformed, and data were normalized based on the average abundance within each sample to account for variation in sampling volumes (Aguilan et al. 2020). Raw data files of the fractions were processed in Proteome Discoverer 3.1 as a single contiguous input file using the same parameters described above. For the PRM analysis, the light-to-heavy ratio between heavy-labeled peptides and endogenous light peptides was determined using Skyline software version 24.1 (MacLean et al. 2010). Transitions were selected based on the intensity ranking. For each peptide, the five most intense fragment ions were monitored.

### EIF4H silencing

HCT116 cells were transfected with siRNAs targeting EIF4H (MISSION esiRNA, Merck) at a final concentration of 20 nM for 48 hours. Control cells were transfected with a non-targeting siRNA (EGFP) under the same conditions. INTERFERin (Polyplus) was used as a transfection reagent.

### Relative cell proliferation

HCT116 cells were seeded at 2 x 10^3^ per well onto 96-well plates and incubated at 37°C in standard conditions. After attaching to the wells, the cells were treated with si-EIF4H or non-targeting siRNA as described previously. The cell number was followed by the high content fluorescent microscope Operetta (PerkinElmer) in digital phase contrast every 24h for 144 h.

### Ribosome stalling assay

HCT116 cells were seeded in a 96-well plate format and transfected with 100 ng of dual reporter stalling constructs (Kriachkov et al. 2023) using the FuGene transfection reagent (Promega). 109-nucleotide long sequences corresponding to a portion of exon 6 of the EIF4H gene and comprising either the Alternative or Reference version of the rs1554710467 tranSNP were cloned on a dual fluorescence plasmid, upstream of an mCherry and downstream of a GFP sequence, separated by P2A sequences which induce the ribosomes to skip the formation of peptide bonds without interrupting translation, allowing the translation of a single mRNA (Lin et al., 2013). A known staller sequence (coding for 17 consecutive lysines) cloned in the same plasmid was used as a positive control. A SEC61B sequence (106 amino acids), which is known to allow read-through, was cloned in the same plasmid and used as a negative control. GFP, mCherry signals, and digital phase contrast were detected after 24-, 36-, and 48-hours post-transfection using the Operetta High-Content Imaging System (Perkin-Elmer). GFP and mCherry intensities were used to calculate GFP/mChFP ratios, which were used as proxies for ribosome stalling. High GFP/mChFP ratios indicate that stalling events occur.

### Image analysis

Images were analyzed using Harmony 4.1 software (Perkin-Elmer). For each fluorescent channel, a Basic Flatfield correction was applied. The DPC channel was preprocessed using the Texture SER method (filter SER ridge, scale 2px, unnormalized) to improve the cell segmentation (Find Cell, Method P). For each cell, the intensity of GFP and mChFP has been measured, and the ratio GFP/mChFP has been calculated. A morphological classification of the cell was applied to exclude round-detaching cells from the healthy ones based on the cellular roundness (<0.8).

### Statistical analysis

Results were plotted and statistically analyzed using GraphPad Prism, version 10. Details on the statistical test applied, the number of replicates, and the significance levels are provided in each figure legend.

## Results

### Using quantitative proteomics of whole lysates to validate missense tranSNPs in HCT116 cells

We leveraged RNA-sequencing experiments we performed on four replicates of total cytoplasmic and polysome-bound mRNA extracts in HCT116 cells cultured in mock conditions, or treated with the MDM2 inhibitor Nutlin (Rizzotto et al. 2020). We also leveraged an equivalent experiment performed on HCT116 cells stably depleted for PCBP2 and DHX30 RBP expression. Applying a recently developed pipeline (Hamadou et al. 2025; Valentini et al. 2021), we identified instances of variation in allelic fraction for heterozygous SNPs and SNVs in comparing total and polysomal extracts. While in a recent report, we focused on instances of imbalances occurring at UTR sequences, here we focused on those that are coding and result in missense changes. We exploited a dataset comprising four biological replicates of three derivative HCT116 cells stably silenced for the RNA-binding proteins DHX30, PCBP2, or control. For each cell clone, total cytoplasmic and polysomal RNAs were extracted from mock or Nutlin-treated cultures and processed for single-end RNA reads. The raw dataset is deposited in GEO and was previously described (Rizzotto et al. 2020). Based on the coverage of the RNA-seq dataset and the filtering applied to define heterozygosity in RNA-seq (AF value between 0.2 and 0.8; AF = Alternative allele read / sum of alternative and reference allele) and coverage (at least 20x) in each biological replicate, 1753 SNP/cell line/condition combinations could be studied for differential allele-specific expression (**Table S1**). To assess the presence of allelic imbalance, for each SNP, we exploited the concordance of differential AF measures across pairs of polysomal and total RNA biological replicates by performing paired t-test statistics. This approach highlighted 120 SNP/cell line/condition combinations (**Table S2**), of which 57 were exonic and comprised 52 unique SNPs/SNVs. Of these, 16 were nonsynonymous tranSNPs showing differential allelic imbalance in RNA-seq that could be interrogated for allele-specific variation in protein expression (**Figure 1A, S1A**; **Table S3**).

**Figure 1.**
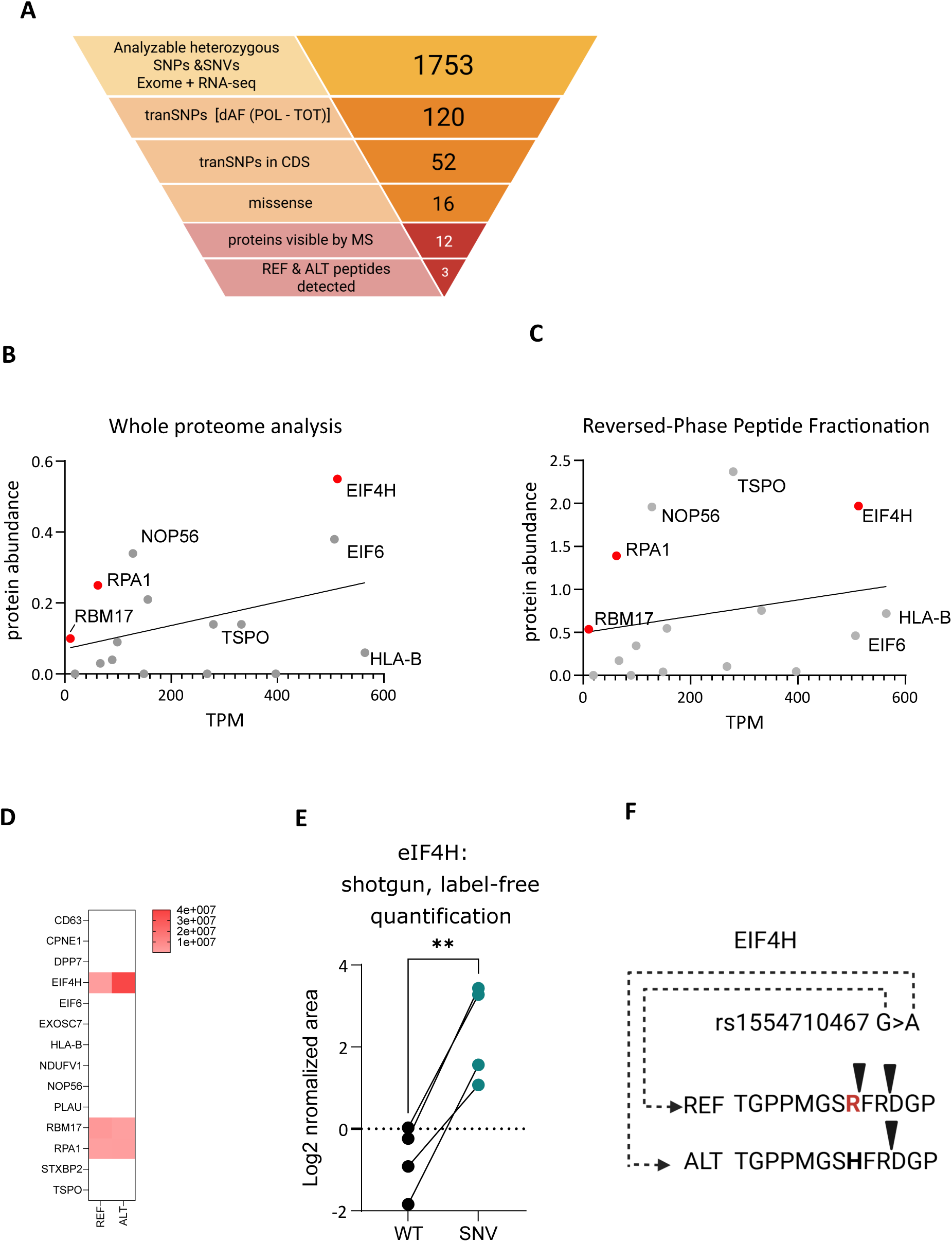
Polysomal profiling and mass-spectrometry identify nonsynonymous allelic variants associated with changes in protein expression. **A)** Workflow of the experimental approach, starting with the number of heterozygous SNPs + SNVs that could be analyzed using HCT116 cells. TranSNPs are defined as those heterozygous variants that showed a significant imbalance in computed allelic fractions in the comparison between polysomal (POL) and total (TOT) RNA sequencing data. To be considered, a minimum threshold of 20 reads for each of the four biological replicates in both fractions was set. The number of tranSNPs present in coding sequences and causing nonsynonymous amino acid changes is indicated. Twelve out of sixteen proteins containing missense tranSNPs were identified by shotgun, label-free whole proteomics, 14 when a peptide fractionation kit was used prior to the analysis. For only three cases, peptides encompassing the heterozygous variants were detected. **B**) Comparison between mRNA expression (TPM, average of the biological replicates) and relative protein abundance measured from whole proteome analysis of HCT116 cells. Highlighted in red are those proteins where the peptides comprising the biallelic variant were detected by mass spectrometry. **C**) Same as B, except that the whole proteome data was obtained after fractionation of digested peptides by High pH Reversed*-*Phase Peptide Fractionation Kit (Thermo Fisher Scientific). **D**) Heatmap presenting the relative abundance of the reference and alternative peptide differing by one amino acid, resulting from the decoding of the heterozygous genetic variants. 14 of the 16 proteins were detected by mass-spectrometry, but for only three of them, the alternative and reference peptides were measurable by whole proteomics. **E**) Relative quantification of the wild-type- and SNV-derived EIF4H peptides by label-free shotgun analysis. ** p< 0.01, two-tailed paired t-test. **F)** The rs1554710467 SNV results in the R183H amino acid change and modifies the cleavage pattern of the two variants, resulting in a 10 amino acid long peptide for the alternative allele. Black triangles indicate the respective cleavage sites for trypsin.

To explore the possibility of using proteomics to quantify the relative protein level being expressed by the two alleles, we first exploited a label-free shotgun proteomics approach and evaluated the correlation between relative mRNA (TPM) and protein abundance. As expected, there was a general positive correlation between those two variables **(Figure 1B)**. Overall, a signal was found for 12 of the 16 proteins containing missense tranSNPs. Given that the protein resulting from the translation of a specific allele can be resolved only by the peptide containing the decoding of the missense SNP or SNVs, protein abundance is an imperfect matrix, given the contribution of additional variables on the possibility of observing a specific peptide from a protein, especially the availability and efficiency of proteolytic sites and enzymes. In fact, after trypsin digestion and LC-MS analysis, a peptide containing at least one of the two biallelic positions was identified in only 3 out of the 12 proteins detected (**Figure 1B**).

Next, to increase proteome coverage, digested peptides were first fractionated using a High pH Reversed-Phase Peptide Fractionation Kit (Thermo Fisher Scientific). The resulting ten fractions were dried by vacuum centrifugation and reconstituted in 0.1% formic acid for LC-MS analysis. This procedure led to a slight increase in the sensitivity of the whole proteome approach, with 14 of the 16 proteins detected (**Figure 1C**). This analysis also enabled the observation of two peptides corresponding to the protein sequence containing the tranSNP for RPA1, RBM17, and, especially, EIF4H (**Figure 1D**). For the remaining 11 proteins, the detected peptides did not overlap with the SNP sites, although for some of them, *e.g.,* NOP56 and TSPO, the abundance was relatively high. Based on these analyses, we selected EIF4H as the most promising candidate, based on transcript and protein abundance and the possibility of detecting peptides corresponding to the translation of the two alleles differing for the rs1554710467 SNV. We also considered RPA1 as an example of a lower-abundance protein.

To further enrich the proteins of interest, HCT116 cell total protein extracts were resolved by SDS polyacrylamide gels electrophoresis, stained with Coomassie, and the gel bands corresponding to the regions of 20-30 kDa (for EIF4H), and 65-85 kDa (for RPA1) were excised and cut into small pieces (∼1 mm^3^). The gel pieces were destained and dehydrated, and in-gel digestion was performed using trypsin (see methods). Digested samples were then processed using shotgun label-free quantification proteomics. Using this approach, significant differences in the abundance of the reference (REF) and alternative (ALT) peptides were observed for EIF4H (**Figure 1E**, **Table S4**). The REF peptide has two potential tryptic sites (8 and 10 AAs long, respectively) (**Figure 1F**).

However, only the 8 AA version of the REF peptide was found in the LC-MS analysis. In the ALT peptide, the amino acid change introduced by the alternative allele generates only a tryptic peptide (10 AAs long), which was found to be more abundant than its REF counterpart. Our finding is also supported by the presence of an aspartic acid (D) adjacent to a tryptic site (R), a sequence context that was reported to inhibit the cleavage efficiency of the enzyme (Šlechtová et al. 2015; Sun et al. 2021). The observed imbalance in the relative abundance of REF and ALT peptides was consistent with the delta allelic fraction measured in the polysomal profiling RNA-seq, showing an imbalance in the polysomal RNA fraction for the alternative allele. By using the same workflow, we investigated the RPA1 protein. However, the results were inconclusive due to the low representation of the reference peptide (not shown).

### rs1554710467 is a coding tranSNP in the EIF4H gene associated with differences in protein allelic variants according to mass-spectrometry analysis

Although present in the dbSNP database, the rs1554710467 is a rare variant, with a reported frequency of 2.75×10^-6^ based on exome sequencing from gnomAD (Chen et al. 2024). EIF4H is a translation initiation factor that, together with EIF4B, can participate in the unwinding of the 5’UTR motif supporting translation initiation (Sun et al. 2012). EIF4H expression is upregulated in some cancer types, and an oncogenic function has been proposed (Vaysse et al. 2015; Wu et al. 2011; Krassnig et al. 2021). Based on our RNA-sequencing data, the alternative allele showed higher allele frequency in the polysomal fraction than the total RNA fraction, inferring that it would lead to higher relative protein expression (**Figure 2A, Table S1**). First, we decided to validate the allelic imbalance by preparing additional biological replicates of polysomal profiling (**Figure 2B**) and using Sanger sequencing as an alternative approach to measuring allelic imbalance. For the EIF4H gene, multiple transcripts are annotated, among which two golden-pass coding ones in the Ensembl database differ in the skipping of exon 5, resulting in a protein shorter by 20 amino acids (**Figure 2C**). rs1554710467 is located in an exon that is common to both splice variants. Hence, we developed specific primers to study each transcript separately (**Table S6**). Allelic fraction calculation by Sanger sequencing was extended to total cytoplasmic RNA, samples corresponding to the density of ribosomal subunits (SUB), the 80S monosomes, light, and heavy polysomes (**Figure 2B, 2D**). For the longer EIF4H isoform, results showed higher minor allele frequency (AF) in the 80S, light polysomes, and a trend in heavy polysomes compared to subpolysomal RNA. For the shorter EIF4H transcript, a trend for higher AF in the 80S was apparent; however, the AF was significantly lower in both light and heavy polysome fractions. Overall, we validated the RNA-seq result showing higher AF for the alternative alleles, but this feature seems to be dependent on the EIF4H transcript that includes exon 5.

**Figure 2.**
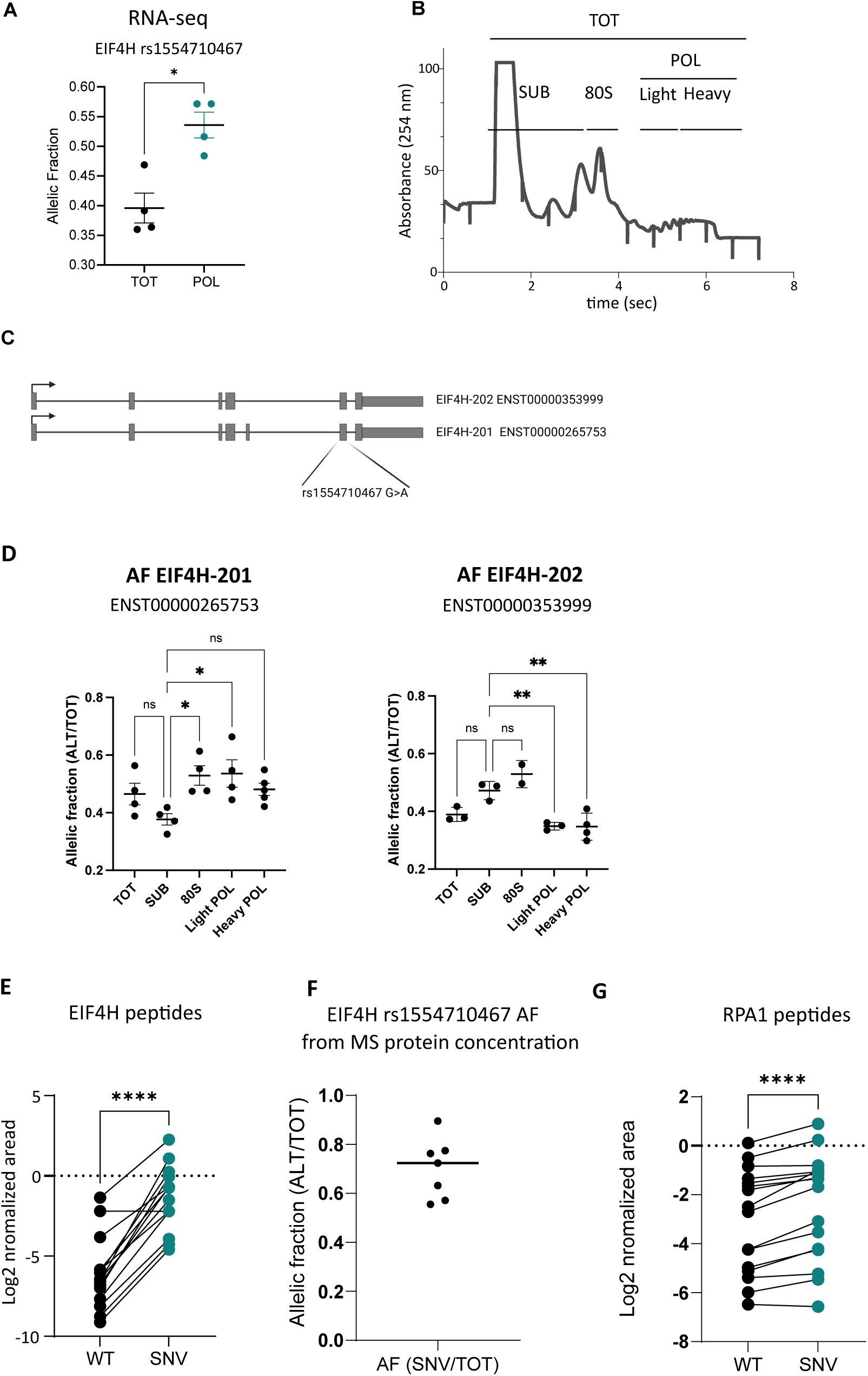
The rs1554710467 is a heterozygous SNV showing consistent allelic imbalance at both mRNA and protein levels. A) Plot comparing allelic fractions (AF, defined as the ratio between the number of reads of the alternative over the total number of reads for that nucleotide position) measured by RNA-seq of total RNA (TOT) and polysome-associated RNA (POL) recovered from sucrose-gradient fractionation. Mean AF, standard deviation, and the individual biological replicates are shown. * p<0.05, two-tailed paired t-test. **B**) Polysome profile of sucrose-gradient density fractionation of HCT116 cytoplasmic lysate. Fractions corresponding to ribonucleoparticles and ribosomal subunits (SUB), 80S monosomes, and light and heavy polysomes were combined as indicated to recover RNA. An aliquot of each fraction was used to reconstitute a cytoplasmic total RNA sample (TOT). **C**) Scheme of EIF4H alternative transcripts. The SNV resulting in the R183H amino acid change is present in an exon that is common to the two annotated coding transcripts. **D**) RNA recovered from polysomal profiling as in B) was used to amplify the SNV exon, followed by Sanger sequencing. Electropherograms were quantified, and results are expressed as AF, calculated as in A). Mean AF, standard deviation, and individual biological replicates are plotted. Specific primers allowed the assay to be performed separately for the two indicated transcript variants, which differ due to an exon skipping event. ns, not significantly different; * p< 0.05; ** p< 0.01, ordinary one-way ANOVA, with Dunnett’s multiple comparisons test. **E**) Relative quantification of the WT and SNV EIF4H peptide from targeted mass-spectrometry analysis employing spiked-in isotope-labeled control peptides. **** p<0.0001, two-tailed paired t-test. The peptide under analysis contains a methionine that can be oxidized during sample preparation. Hence, the plot shows paired results for both the unoxidized and oxidized peptides. **F**) Quantification of AF, defined as in A) from multiple mass-spectrometry experiments. Median and individual results are shown. The SNV (ALT) peptide is more abundant than the WT (REF). **G**) Relative quantification of the WT and SNV RPA1 peptide by targeted mass-spectrometry performed as in E). See Table S1 for AF data of the nonsynonymous rs5030755 variant. Although highly significant (**** p<0.0001, two-tailed paired t-test), the difference in the relative abundance of the peptides derived from the two alleles is smaller compared to what we observed with EIF4H.

After confirming the allelic imbalance within ribosomes, we returned to a mass-spectrometry approach to explore whether the two alleles could be quantified.

To this aim, a targeted proteomics approach was chosen. Heavy isotope-labeled synthetic peptides representing REF and ALT variants were spiked into the endogenous peptide mixture to increase the confidence of identification and quantification in LC-MS analysis. The results of MS-based targeted proteomics confirmed the significant increase in the abundance of the alternative rs1554710467 protein allele (**Figure 2E**), as predicted by polysomal profiling. The relative abundance of the two peptides in the samples was estimated by the ratio of the endogenous (light) peptide to its spiked (heavy) synthetic counterpart, using the Skyline software to compute the peptide allelic fraction (**Figure 2F**).

Overall, the mass spectrometry results confirmed an allelic imbalance for the EIF4H rs1554710467 that was even higher than what was measured by polysomal profiling, RNA-seq, or Sanger sequencing. The targeted approach and the use of isotope-labeled peptides enabled us to validate the imbalance also for RPA1 rs5030755-spanning peptides (**Figure 2G**).

### The rs1554710467 variant does not affect the EIF4H protein half-life or its subcellular localization

We wanted to exclude the possibility that the difference in relative abundance of the two EIF4H peptides could depend on the effect of the amino acid substitution (R183H) on protein stability. To gain information in this respect, HCT116 cells were treated with cycloheximide, and the EIF4H protein half-life was measured in a time-course experiment. Results established that EIF4H is a relatively long-lived protein with an estimated half-life of about eight hours in our experimental conditions (**Figure 3A**). We then prepared protein extracts at time 0 and after 8 hours of cycloheximide treatment and performed again the targeted mass spectrometry analysis. Results established an equivalent imbalance of the two peptides corresponding to the rs1554710467 alleles, suggesting that the two alternative peptides belong to proteins possessing similar half-lives (**Figure 3C**).

**Figure 3.**
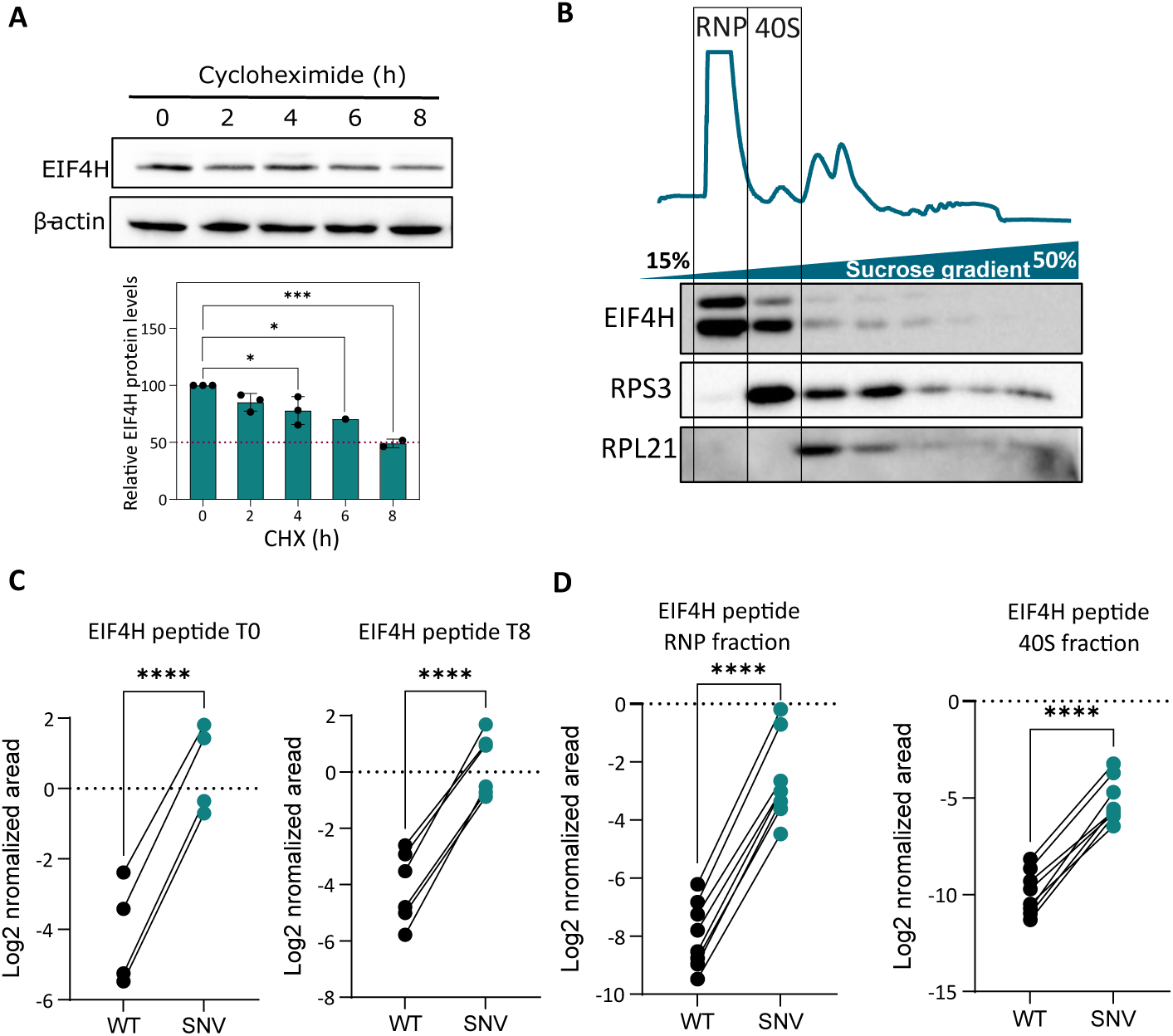
The rs1554710467 SNV does not alter EIF4H protein stability nor its subcellular localization. **A**) A cycloheximide chase assay was performed to estimate EIF4H half-life in HCT116 cells. Top panel, representative western blot for EIF4H and beta-actin proteins. Bottom panel, relative EIF4H quantification based on densitometric analysis of the immunoblots. The average relative signal, the standard deviations, and the individual replicates are shown. * p< 0.05; *** p< 0.001, ordinary one-way ANOVA. **B**) HCT116 polysomal profiling was used to extract protein from each fraction and perform western blot analysis for EIF4H, RPS3, and RPL21 proteins. Based on these results, proteins recovered from the RNP and 40S fractions were used for targeted proteomics. **C**) Based on cycloheximide chase assay results, proteins at time 0 and at the 8-hour time point (corresponding to half-life) were subjected to targeted proteomics. D) Based on the EIF4H protein localization obtained in B), proteins recovered from the RNP and 40S fractions were used for targeted proteomics. **** p<0.0001, two-tailed paired t-test.

Next, we explored whether the EIF4H alternative-allele protein could retain its function. To support this aim, proteins were retrieved after sucrose gradient fractionation of HCT116 cytosolic extracts. Western blot analysis revealed that the EIF4H protein was mainly localized with the light fraction corresponding to RNPs and with the fraction corresponding to the 40S subunits (**Figure 3B**). Proteins recovered from these two fractions were subjected to the targeted mass spectrometry analysis, which showed again a significant imbalance between the two alleles (**Figure 3D**). We concluded that the amino acid change associated with rs1554710467 seems to preserve protein half-life and subcellular localization, suggesting that the alternative allele is functional. Furthermore, protein structure predictions by AlphaFold 3 (Abramson et al. 2024) suggest that the R183 amino acid is part of a relatively long unstructured domain, and amino acid changes at that residue are predicted to be benign **(Figure S1B, S1C)**. Overall, we suggest that R183H represents a gain-of-function allele due to higher mRNA translation potential.

### The alternative rs1554710467 allele shows higher translation in a ribosome stalling assay

To investigate whether the amino acid change caused by the rs1554710467 could affect translation efficiency by some type of ribosome stalling, we cloned 109-nt sequences of EIF4H exon 6, comprising either the alternative or the reference allele of the rs1554710467 tranSNP, on a dual-fluorescence plasmid, between a GFP and an mCherry sequence and flanked by two P2A sequences (Kriachkov et al. 2023). The SEC61B sequence, which is known to allow efficient read-through, was cloned in the same plasmid and used as a negative control. A positive control for stalling, containing a sequence coding for a stretch of seventeen consecutive lysine residues, was used (**Figure 4A**). With these reporter constructs, the GFP/mChFP can be used as a proxy for ribosome stalling. The two EIF4H alleles were similar to the SEC61B control, although being about one-third in length, highlighting that the context around the rs1554710467 tranSNP is not particularly complex for the translation machinery. However, while the GFP/mChFP ratio for the reference EIF4H G allele was not significantly different compared to the control, the alternative A allele produced a significantly lower GFP/mChFP ratio, indicating that it is translated more efficiently (**Figure 4B, Figure S3)**. This difference is consistent with both the allelic imbalance in RNA-seq and the mass spectrometry experiments, with all three assays pointing to higher EIF4H protein levels from the alternative allele (**Figure 4B**, **Figure S3B**). The higher translation efficiency for the alternative allele was also confirmed at different time points after transfection of the dual fluorescent reporter plasmid (**Figure S3**). The positive control for ribosome stalling led to a high GFP/mChFP signal ratio as expected (**Figure S3**).

**Figure 4.**
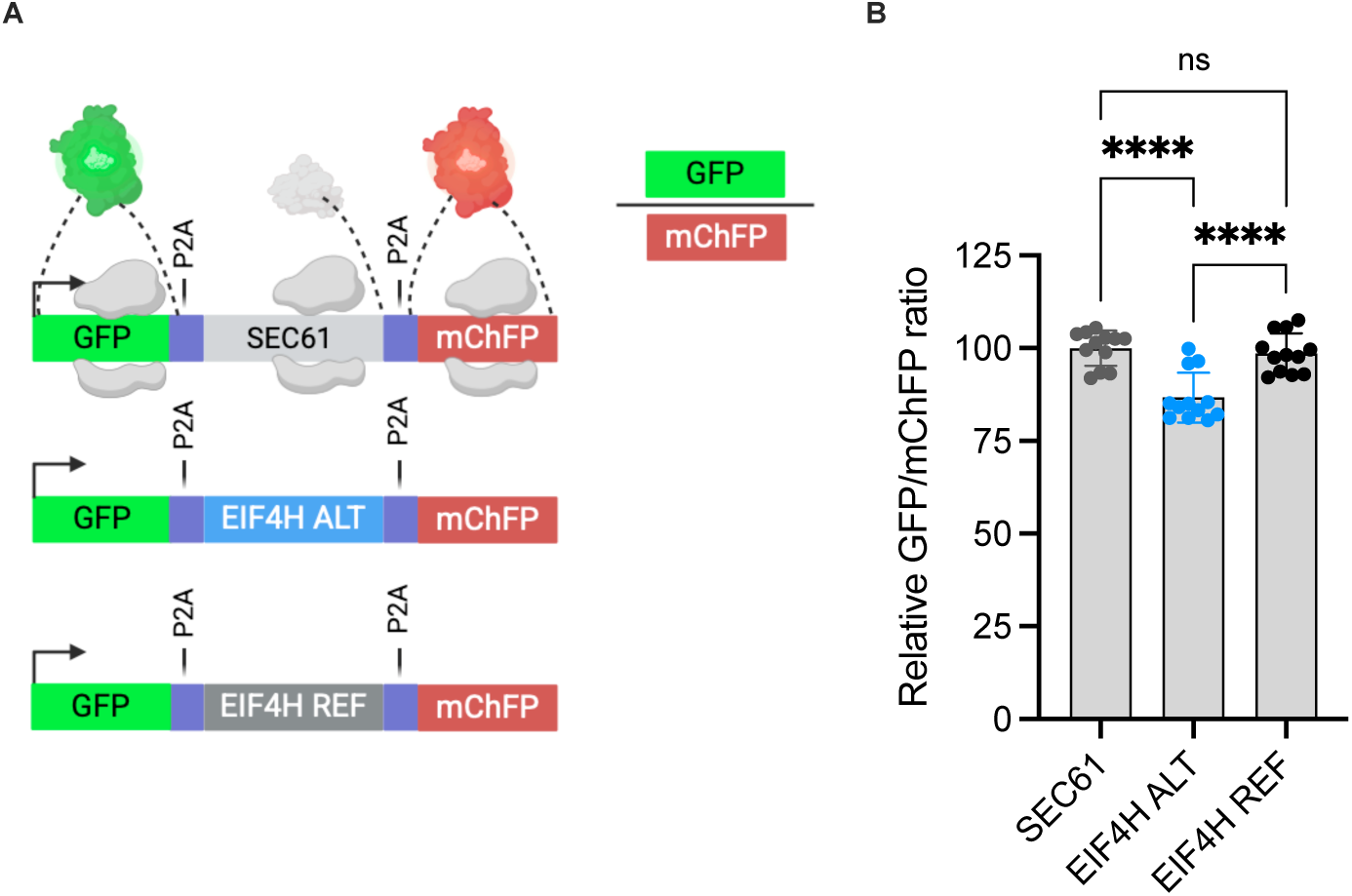
The *rs1554710467* variant shows higher translation in a ribosome stalling assay. A) Scheme of the dual-fluorescence transcript used to evaluate the relative efficiency of translation of a 33 amino acid region of EIF4H exon 6 encompassing the SNV position. The two separated alleles were cloned as intervening sequences between a GFP coding sequence lacking a stop codon and the mChFP coding sequence. As indicated, the intervening cloned sequence is flanked by P2A sites. This configuration enables the production of GFP and mChFP by the decoding of a single transcript. Hence, the GFP/mChFP ratio can be used to measure differences in the efficiency of translation of the cloned sequence. **B**) Relative GFP/mChFP fluorescence ratio measured by Operetta, HTS microscope. Average, the standard deviations and the individual ratio of independent replicates are shown, setting to 100% the results obtained with the SEC61 sequence, used as a control of a sequence not associated with ribosome stalling. ns, not significantly different; **** p< 0.0001, ordinary one-way ANOVA, with Sidak’s multiple comparisons test. At least 1000 cells were segmented and quantified for each replicate. The distribution of fluorescence ratio per cell is presented in Figure S2.

### Depletion of EIF4H does not overtly impact cell proliferation but is associated with reduced RPS6 protein expression

To explore the potential functional consequences of the rs1554710467 resulting in higher EIF4H expression, we attempted to partially deplete EIF4H in HCT116 cells by transient siRNA delivery. Although this approach was not allele-specific, it was performed to explore the consequences for the cells of a reduction in EIF4H expression, to counteract the proposed effect of the SNV. Knockdown efficiency was confirmed by qPCR and Western blot analysis 48 hours post-transfection (**Figure 5A-5C**). Based on EIF4H’s known function in unwinding structured 5’UTR motifs during translation initiation, we hypothesized that reducing EIF4H levels could specifically impair the translation of TOP mRNAs and tested RPS6. EIF4H depletion was associated with reduced RPS6 levels (**Figure 5B-5D**). Instead, EIF4H depletion was not associated with a reduction in cell proliferation (**Figure 5E**).

**Figure 5.**
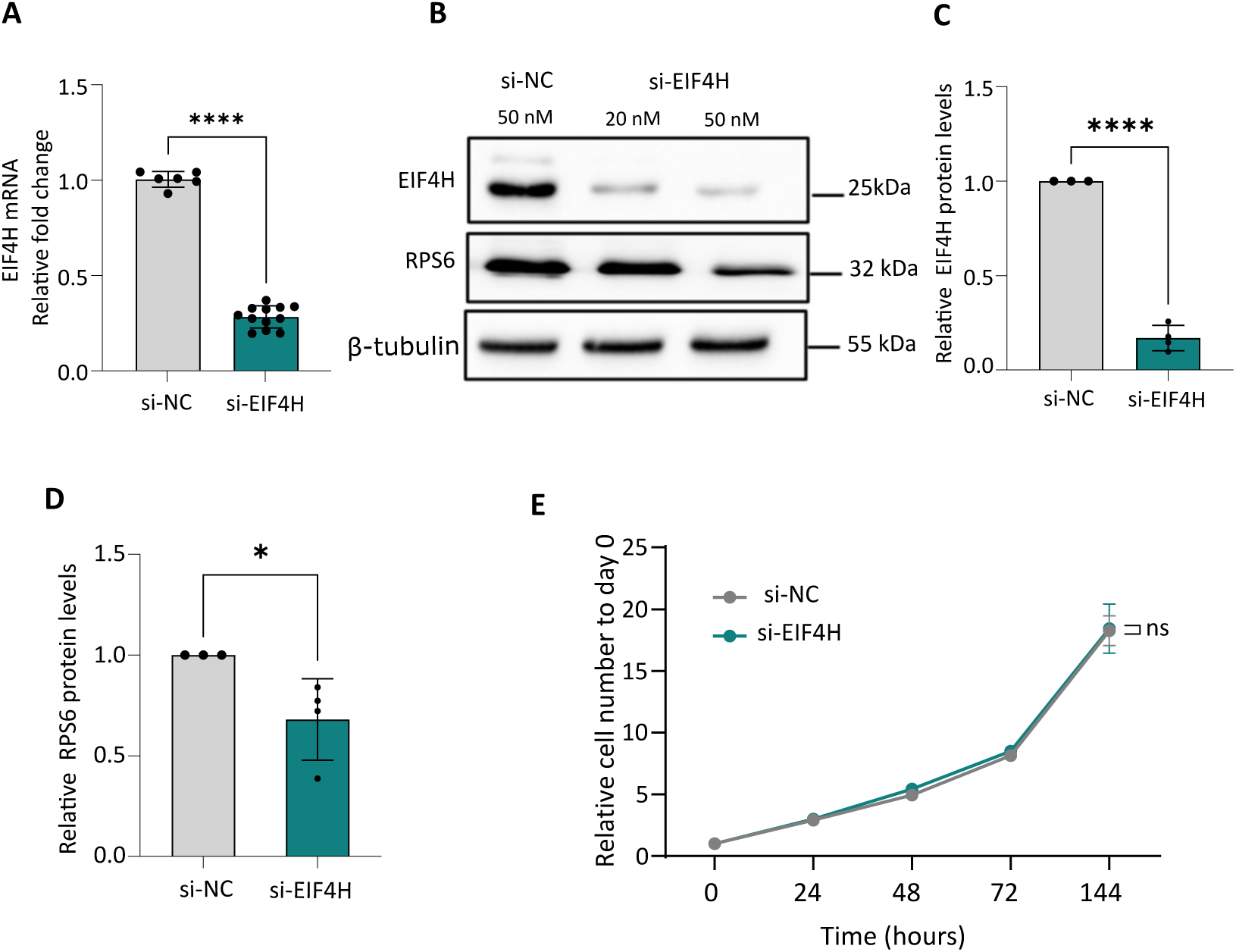
EIF4H depletion in HCT116 cells does not impact proliferation but results in lower expression of RPS6 protein. **A)** Relative EIF4H mRNA expression in HCT116 cells transfected with a non-targeting siRNA control or siRNAs targeting EIF4H. The average relative expression, standard deviation, and individual replicates are shown. **** p<0.0001, two-tailed unpaired t-test. **B**) Representative western blot showing the reduction of EIF4H protein in HCT116 cells 72 hours after the transfection of siRNAs at the indicated concentrations. RPS6 protein was also detected. Beta-actin was used as a reference protein. **C**) Efficiency of EIF4H silencing at the protein level based on the quantification of western blot images. **** p<0.0001, two-tailed unpaired t-test. **D**) Impact of EIF4H depletion on the relative expression of RPS6. * p<0.05, two-tailed unpaired t-test. **E**) Relative proliferation of HCT116 cells transfected with a non-targeting siRNA control or siRNAs targeting EIF4H. Cell number was measured by digital phase contrast using Operetta. Results are expressed relative to the T0 cell number. ns = not significant, 2-way ANOVA.

## Discussion

A broad change in cellular metabolism is one of the established hallmarks of cancer cells, as exemplified by altered expression or acquired somatic mutations leading to looser control of cell growth and the balance between proliferation, autophagy, and cell death patterns. Along with the growing evidence for unexpected complexity and variability in ribosome composition and function, these findings justify the growing interest in studying mRNA translation control in cancer and the opportunity for developing effective cancer therapies targeting translation initiation (Bhat et al. 2015). Importantly, there is considerable evidence that transcripts coding for elements of the translation machinery in a broad sense are themselves subject to translational control mechanisms, such as regulation of translation initiation, including cap-independent translation or the interplay between overlapping open reading frames (Ingolia et al. 2009; Hsieh et al. 2012; Thoreen et al. 2012; Meyuhas 2000).

Along these lines, we are developing methods to annotate genetic variants associated with alterations in gene expression that appear to be linked to changes in mRNA fate in the cytoplasm. For example, our earlier studies revealed that the commitment to p53-dependent apoptosis can be modulated by an mRNA translation program comprising the RNA helicase DHX30, the RNA binding protein PCBP2 acting at a target cis-element present in the 3’UTR of pro-apoptotic mRNAs (Rizzotto et al. 2020; Zaccara et al. 2014). This finding stimulated us to annotate genetic sources of allele-specific mRNA translation potential in cancer-relevant genes that were sought out by exploiting RNA-seq data of matched polysomal and total RNA (Valentini et al. 2021). The underlying assumption is that the translatome is not only a good proxy of the proteome at the gene level, as broadly supported in the literature (Schwanhüusser et al. 2011; Ingolia et al. 2009; Battle et al. 2015), but also at the allelic level when the two alleles are expressed from a heterozygous cell line.

A consistent variation in the allelic fractions in the comparison between polysomal and total RNA for heterozygous SNPs is the initial criterion we use to nominate a tranSNP, defined as a genetic variant associated with allele-specific mRNA translation. Using polysomal profiling, we had the opportunity to also study heterozygous SNPs and SNVs present in the untranslated regions and pursue underlying mechanisms. Indeed, UTR tranSNP could modify cis-elements for microRNAs or RBP binding, which in turn can impact the transcript fate in terms of subcellular localization, translation efficiency, or stability.

In this study, we established the feasibility of using allele-specific proteomics to uncover instances of variation in translation efficiency among nonsynonymous coding SNP and SNV alleles present in the heterozygous state in the HCT116 cancer cell line. Compared to translatome and RNA sequencing, the sensitivity of MS approaches is still limited, and the detection of SNV and SNP-containing peptides can be hampered by their low abundance and low protein coverage due to a lack of enzymatic cleavage sites. In order to improve the monitoring of SNV-containing peptides, we have combined different strategies using fractionation and shotgun proteomics analysis as a first step, followed by targeted PRM proteomics to achieve superior quantitative accuracy (Figure 1). By using these methods, we were able to develop a proof-of-principle study for EIF4H and the rs1554710467 missense SNV. Our results consistently showed a pronounced difference in the relative abundance of the two allelic peptides produced by EIF4H protein digestion. The imbalance was consistent with the changes in allelic fraction from RNA-seq, validated by Sanger sequencing. A consistent and significant difference in the mRNA translation potential was also measured by dual fluorescence assays originally developed to study ribosome stalling. Interestingly, the magnitude of the effect of the single R>H amino acid variant we studied is equivalent to the impact observed with 10 copies of GR repeats (Kriachkov et al. 2023). We provided evidence that the imbalance seen by mass spectrometry cannot be explained by differences in protein stability among the two alleles (Figure 3). Also, results based on protein extracts from sucrose-gradient fractionations suggest that the wild-type and variant proteins have equivalent subcellular localization (Figure 3). Structural modeling does not predict that the R183H change has an overt impact on protein folding. However, we cannot exclude that the histidine change will lead to a more tightly structured motif elicited by EIF4H interactions with a target protein. Overall, we propose that the rs1554710467 SNV represents a gain-of-function allele as it leads to higher levels of an EIF4H protein that is predicted to be functional.

Several mass spectrometry-based approaches have been developed to empower the detection and quantification of protein biomarkers, especially cancer-associated mutant proteins, in complex biological samples, as well as in cancer cell lines (Halvey et al. 2012; Ogata et al. 2022; Wang et al. 2011; Lin et al. 2023; Tan et al. 2020; Shi et al. 2018; Tan et al. 2017; He et al. 2020; Johansson et al. 2013). However, there are limitations in MS-based techniques such as selected reaction monitoring (SRM), multiple reaction monitoring (MRM), and parallel reaction monitoring (PRM) (Wu et al. 2013; Wu and Snyder 2015; Ogata et al. 2022). In fact, global proteomics strategies typically detect only a fraction of single amino acid variants identified via DNA- or RNA-based sequencing due to issues like low peptide abundance, lack of matching peptide sequences in databases, and relatively high false discovery rates (Wu et al. 2013; Wu and Snyder 2015; Ogata et al. 2022; Lin et al. 2023; Sheynkman et al. 2014).

Our proof-of-principle experiments aimed at integrating transcriptome, translatome, and mass-spectrometry data, leveraging SNPs and SNVs present in the exome that, by being heterozygous, enabled us to quantify allele frequencies both in total RNA and polysome-associated fractions. Hence, the analysis of the variation in allele frequency between polysome-associated and total RNA allowed us to candidate nonsynonymous variants as potentially associated with allele-specific translational regulation (Figure 1A, S1A, Table S1, S2). This pre-selection and the fact that two alleles are expressed by the same cell strongly reduced the number of instances that could be validated, but at the same time, empowered our mass-spectrometry approach, reducing sample-dependent variables that are unavoidable when comparing different cell extracts or, even more so, biological fluids. For example, a large-scale mass spectrometry-based approach led to the detection of over 400 peptides with single amino acid variants. Matched RNA-seq did not support the presence of heterozygous alleles, suggesting a high number of false positives. The correspondence with a biallelic state improved markedly when the analysis was restricted to high-abundance transcripts. For those cases (∼50), allele-specific peptide expression showed greater variability than allele-specific RNA expression, a difference that was attributed to experimental variability in mass spectrometry (Sheynkman et al. 2014). Based on the results of our pipeline, we propose that a matched transcriptome and translatome RNA-seq experiment would have revealed a fraction of those instances of allele-specific peptide expression, as evidence of cis-based mRNA post-transcriptional regulation. A more recent study used an elegant system to produce standards for pairs of reference and alternative peptides to quantify the ratio between peptides derived from wild-type and variant alleles across different heterozygous samples to investigate the possibility of cis-regulatory variants contributing to allele-specific expression within a population (Shi et al. 2018). That study also revealed no significant correlations between allele-specific variations at peptide or mRNA levels for the gene being analyzed, UGT2B15, a finding interpreted as evidence that cis-acting variants regulating the mRNA fate, including translation, were at play. Our approach, particularly using polysomal profiling to obtain translatome data, is tailored to studying genetic sources of mRNA post-transcriptional regulation. In fact, the availability of haplotype data could enable the exploitation of non-synonymous missense variants quantifiable in mass spectrometry as markers of allele-specific protein expression that could be caused, for example, by SNP alleles in UTRs altering translation efficiency, for example, by changing RBP binding sites.

In the case of EIF4H, no other SNPs are present in HCT116, hence the proposal that the rs1554710467 SNV is causative of the observed differences in protein expression. EIF4H is an RNA helicase involved in translation initiation, in particular of transcripts featuring highly structured 5’-UTR, including cancer-relevant ones such as c-MYC, Bcl-xL, and FGF2, possibly modulating cell proliferation, survival, or angiogenesis (Vaysse et al. 2015). While considered as an alternative partner to EIF4A competing with EIF4B, recent structural studies suggested the possibility that both EIF4H and EIF4B can be simultaneously present at translation initiation complexes and also participate in the process of mRNA decoding beyond initiation (Brito Querido et al. 2024). It has been proposed that EIF4H can be considered an oncogene in some cancer types (Vaysse et al. 2015; Wu et al. 2011; Krassnig et al. 2021). Hence, our results suggest that the rs1554710467 SNV could be considered as a gain-of-function allele, as it leads to higher translation potential. However, while highly exploratory, our EIF4H depletion experiment did not reveal an overt dependency of HCT116 cells on high EIF4H levels, although we saw a reduction in the steady-state expression of RPS6, considered a TOP mRNA (Jia et al. 2021), which can be consistent with the canonical role of the EIF4H helicase.

To validate differences in allelic fractions between total and polysomal RNA, we have used surrogate assays, such as Sanger sequencing, targeted resequencing, or reporter gene assays with cloned transcript portions of each of the two allele pairs (Hamadou et al. 2025; Valentini et al. 2021), and for the first time in this study, allele-specific proteomics. In principle, for those heterozygous tranSNPs present in the coding sequence, Ribo-seq data could also be used to measure allelic imbalance, comparing RNA-seq with ribosome-protected fragments. To explore this possibility, we inspected published datasets of HCT116 cells, starting from the GWIPS database (Michel et al. 2014) and retrieving data visualized and analyzed by exploiting the Integrated Genome Viewer (Figure S3, Table S7). While limited by low coverage, we evaluated 8 nonsynonymous and 10 synonymous tranSNPs from our list. As a comparison, we used 20 SNPs showing no evidence of imbalance in our dataset. Interestingly, we observed a difference in absolute delta allelic fraction comparing Ribo-seq with RNA-seq data. The difference was more evident when only non-synonymous tranSNPs were considered. The two groups (tranSNPs and control SNPs) were not different regarding coverage, both for the RNA-seq and Ribo-seq (**Figure S3D**). This preliminary analysis suggests that Ribo-seq datasets could be mined to search for allele-specific differences in ribosome footprints, although low coverage represents a limitation.

While the difference between the two rs1554710467 in terms of polysome-association, translation stalling, and protein expression was very consistent (Figure 2), the mechanisms by which the G>A nucleotide change in the mRNA or the resulting R183H amino acid change in the protein leads to higher translation potential remain to be established. RNA structure predictions by MutaRNA (Miladi et al. 2021) did not show a profound effect of the SNV, considering the two transcripts that differ in the inclusion of exon 5 (**Figure S4**). Mining of eClip data indicates that the SNV site overlaps with binding sites for RPS3 and G3BP1 (not shown), but the significance of these results remains to be established. At the protein level, the sequence surrounding the SNV site is surrounded by two PPX motifs, which can result in slower translation or even ribosome stalling (Gutierrez et al. 2013; Schuller et al. 2017; Han et al. 2020), and the reference allele codes for the first arginine of an RFR motif. Although these features may indicate an impact on the efficiency of decoding that particular sequence context, further studies are needed to dissect the exact mechanism.

In conclusion, we used different mass-spectrometry methods based on whole proteome, fractionation, and targeted analyses with spiked-in isotype-labeled controls to quantify allele-specific differences in protein expression associated with non-synonymous genetic variants (**Figure 6**). We validated this novel approach, although it showed lower sensitivity compared to the translatome RNA-seq-based approach, which limited the analysis to only abundant proteins. In fact, the need to quantify a specific peptide, harboring the bi-allelic marker, within a protein represents an additional limitation compared to other allele-specific proteomics approaches (Yao et al. 2018; He et al. 2020; Johansson et al. 2013; Wu et al. 2013; Shi et al. 2018; Tan et al. 2017).

**Figure 6.**
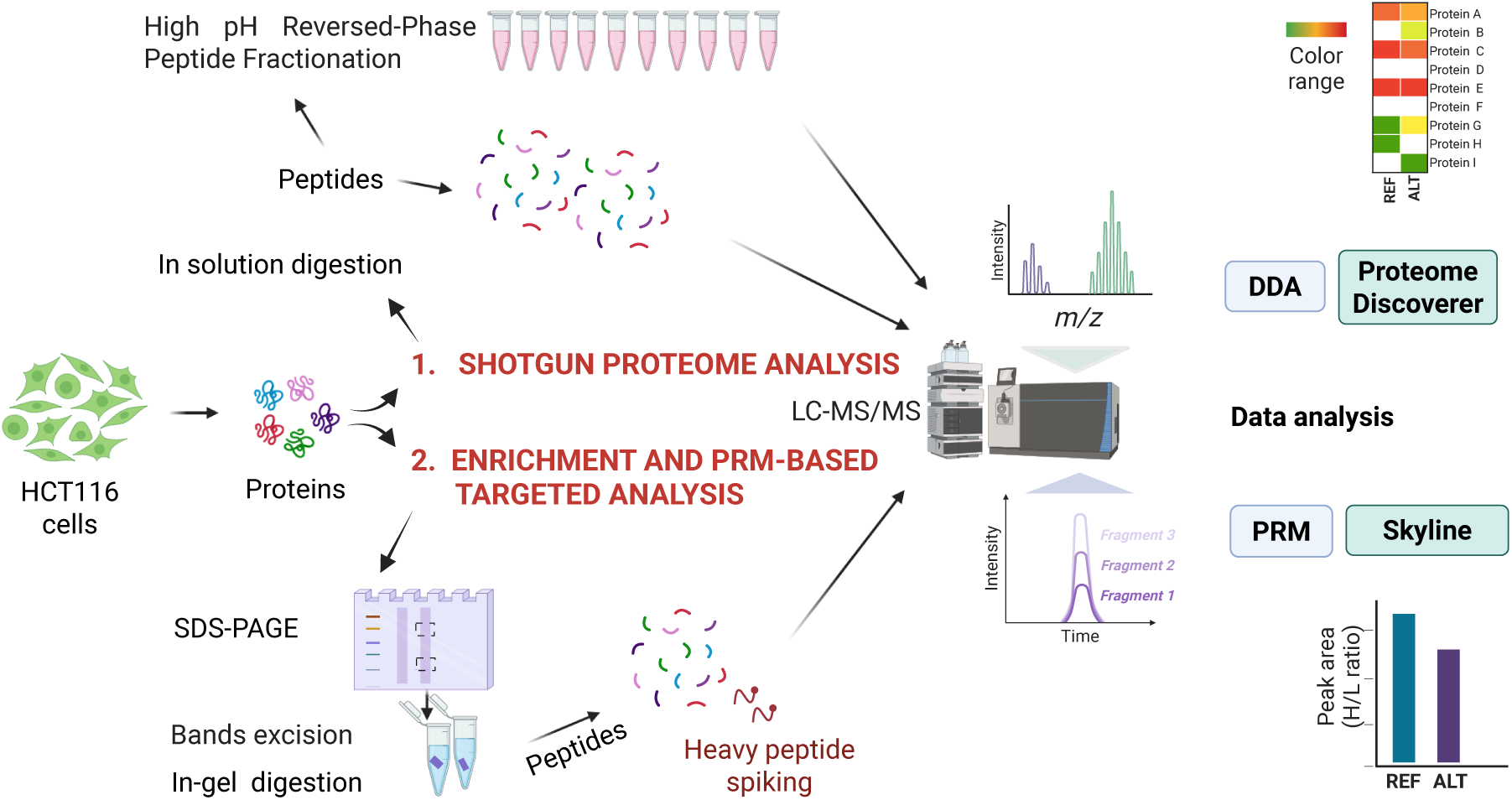
Schematic representation of the allele-specific proteomic approaches used in this study. For the shotgun analysis (**1.**), before the detection, digested peptides were also separated into 10 fractions using a High pH Reversed-Phase Kit. Compared to the unfractionated peptide preparations, this approach slightly increased the number of proteins containing nonsynonymous variants that were detected. However, out of 16 proteins that were followed based on the polysomal profiling RNA-seq data, 14 were detected by the presence of at least one unambiguous peptide, but only in three cases, a peptide encompassing the target variant was revealed. To increase the sensitivity, a targeted proteomics approach was used (**2.**) that relied on protein isolation after acrylamide electrophoresis, in-gel digestion, and spike-in of isotope-labeled peptides. This approach enabled the quantification of the reference and alternative EIF4H peptides from total protein extracts and proteins recovered from sucrose gradient fractions corresponding to the density of light ribonucleoparticles and 40S ribosomal subunits.

This limitation could be partially reduced by using multiple proteases to increase the chance of detecting variant peptides (Sheynkman et al. 2014). Instead, the possibility of studying biallelic markers from the same cell extract guided by translatome data is a strength of our approach, as it reduces experimental and biological variability compared to proteomic approaches to detect somatic mutations, such as cancer-specific biomarkers (Ogata et al. 2022; Wang et al. 2011; Lin et al. 2023).

## Acknowledgments

We thank Massimo Andreis for technical support during his Bachelor’s internships. We thank Prof. Alessandro Provenzani and Dr. Elisa Facen, Department CIBIO, University of Trento, for support with the ribosome stalling reporter assay. We also thank the CIBIO Proteomics and MS core facility at the University of Trento for supporting the experiment optimization. We thank Drs. Gabriella Viero and Gaurav Sharma, Institute of Biophysics, CNR Unit at Trento, Italy, for discussions and support with the sucrose gradient fractionation. We thank Massimiliano Clamer, CEO of Immagina Biotechnologies Srl, for stimulating discussions.

## Supplementary Figures and Tables

**Figure S1.**
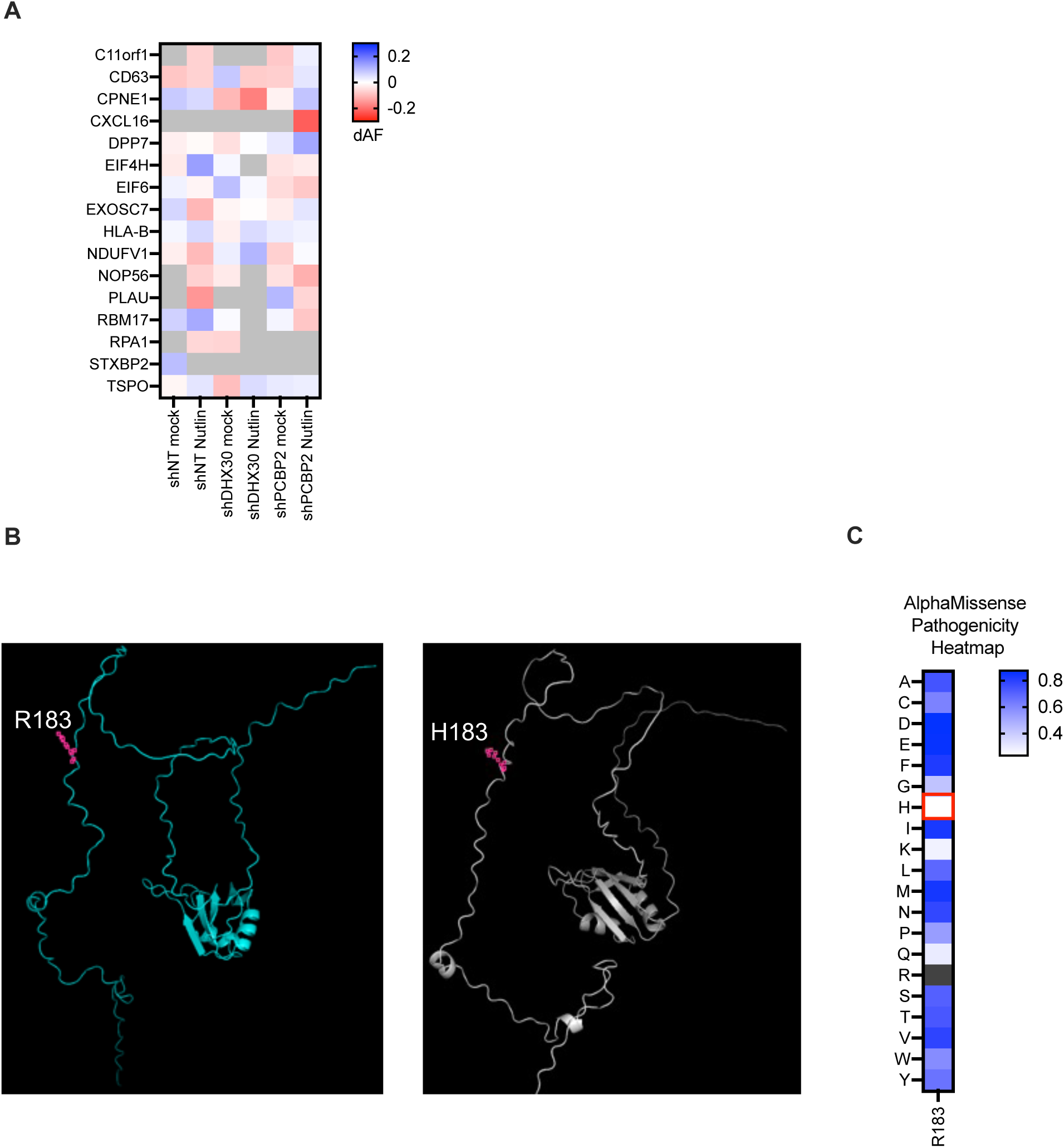
The *rs1554710467* variant leads to the R183H change in EIF4H that is predicted to reside in an unstructured region and to represent a benign amino acid change. A) Heatmap view of variation in allelic fractions (dAF) in the comparison between polysomal and total RNA for 16 nonsynonymous tranSNPs identified as significant in this study. The list of all the heterozygous SNPs and SNVs that could be interrogated in our dataset can be found in Table S1. The RNA-seq raw data is deposited in GEO (GSE95024). RNA-sequencing was performed on control HCT116 cells and on derivative clones stably depleted for DHX30 and PCBP2 expression, two RNA-binding proteins implicated in p53-dependent apoptosis (Rizzotto et al. 2020). **B**) AlphaFold3 predictions of EIF4H R183 and H183 protein structures. The position of the 183^rd^ amino acid is indicated in purple. **C**) Heatmap of the predicted functional impact of amino acid changes at the EIF4H R183 position, according to AlphaMissense. The R183H variant (red square) is classified as “likely benign”.

**Figure S2.**
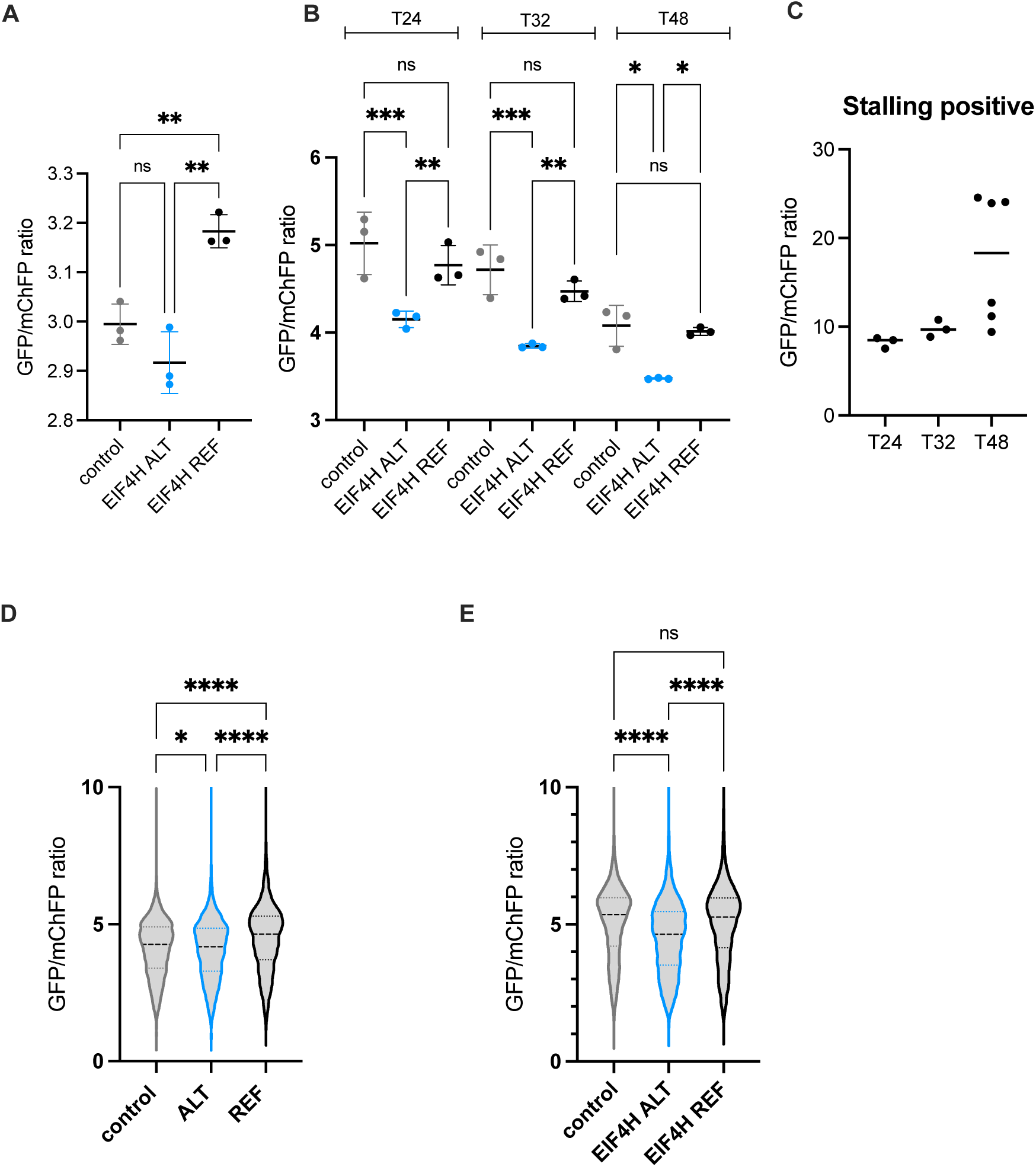
GFP/mChFP ratio obtained with the ribosome stalling reporter plasmids -supporting the data presented in. Figure 4**-. A**), **B**), average GFP/mChFP signals ratio for two independent experiments, each consisting of three independent transfections. For the experiments whose results are plotted in B, analysis was performed at three different time points after transfection. ns, not significantly different; * p< 0.05; ** p< 0.01; *** p< 0.001, ordinary one-way ANOVA, with Sidak’s multiple comparisons test. **C**) GFP/mChFP signals ratio obtained with a ribosome stalling control consisting of a poly-lysine sequence stretch cloned in-between the GFP and mChFP CDS. The average ratio and the data for individual transfected wells are shown. In this assay, a higher ratio signals lower mChFP translation. **D**) **E**) Violin plots of the GFP/mChFP ratio for the two independent experiments. In both cases, numbers were acquired by Operetta HTS 48 hours after transfection. ns, not significantly different; * p< 0.05; **** p< 0.001, ordinary one-way ANOVA, with Sidak’s multiple comparisons test.

**Figure S3.**
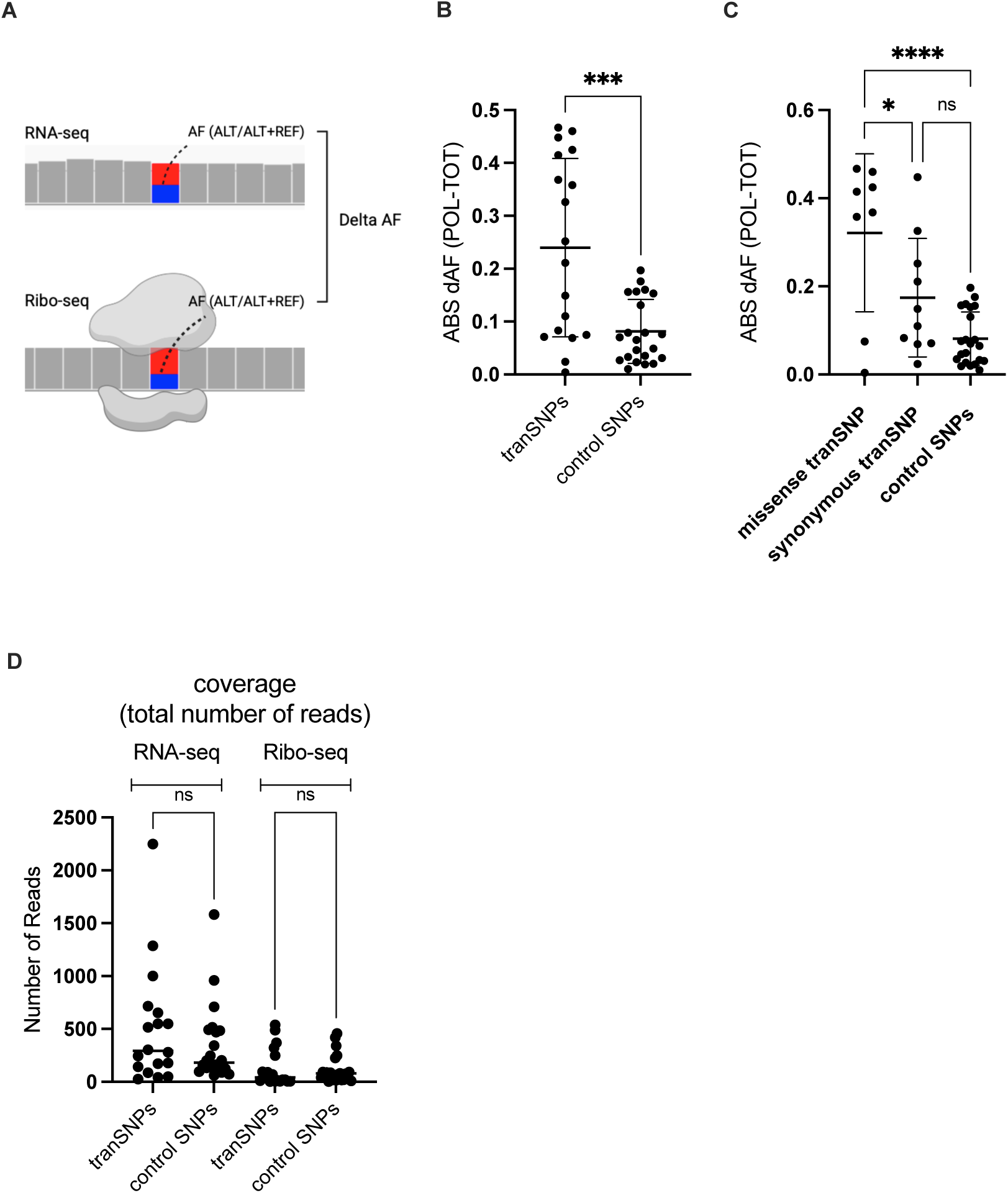
Ribo-seq data can be used to measure allelic fractions and support the instance of allelic imbalance observed by polysomal profiling. **A)** Scheme of the approach. RNA-seq and Ribo-seq data were retrieved starting from the GWIPs database (SRR1333393; SRR4293696; SRR4293698). Coverage and allelic fraction (AF) were measured from the BAM files in IGV(Robinson et al. 2011). Delta allelic fractions were calculated by subtracting the AF measured in Ribo-seq (treated as corresponding to the polysomes in our approach) and in RNA-seq (total RNA). Note that in Ribo-seq data, 80S monosomes are not resolved from polysomes. **B**), **C**) Comparison of absolute delta allelic fractions for SNPs showing allelic imbalance in our polysomal profiling and a control group of SNPs that show similar AF across RNA fractions. Unlike the case of the mass-spectrometry approach, Ribo-seq data allows us to explore all coding variants (nonsynonymous and synonymous). **D**) Comparison of RNA and Ribo-seq coverage (number of reads) for the tranSNP and control group.

**Figure S4.**
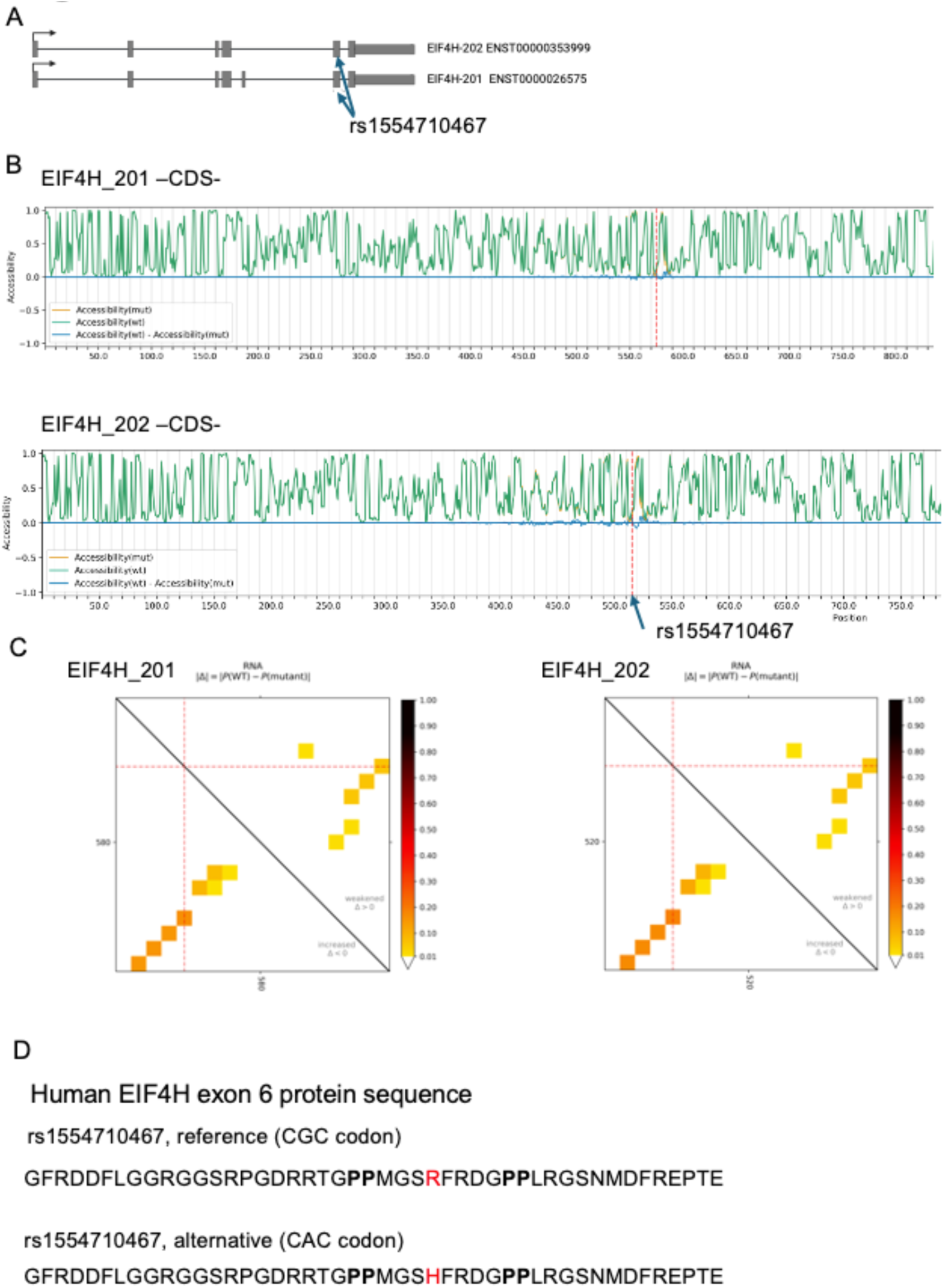
The *rs1554710467* variant is not predicted to alter dramatically the EIF4H mRNA structure. **A**) Scheme of the EIF4H coding transcripts that differ for the exon 5 skipping event. **B**) MutaRNA-derived predictions of relative structural changes for the reference and alternative alleles. The image shows the predictions for the coding sequences, although in the modeling, the 3’UTR sequence up to the limit of sequence length for the software was included. **C**) Blots from MutaRNA highlighting predicted changes in RNA folding energy and base pairing potential between reference and alternative alleles. **D**) EIF4H exon 6 protein sequence. In bold, two PPx motifs. The reference and alternative alleles are indicated in red.

## List of Supplementary Tables

**Table S1** List of analyzable heterozygous SNPs in HCT116 (shNT, shDHX30, shPCBP2, mock; Nutlin) RNA-seq dataset.

**Table S2** List of the tranSNPs identified. ID, gene name, location, and AF data are reported.

**Table S3** List of the synonymous and nonsynonymous coding tranSNPs identified. ID, chromosomal coordinates, gene name, gene and transcript ID, cDNA & CDS position, amino acid change, and impact on protein sequence are reported.

**Table S4A** Selected ion masses and charge states for the ion prioritization method.

**Table S4B** Isolation list for the heavy and light peptides with mass list and charge state information

**Table S5** List of the antibodies used in this study.

**Table S6** List of primers used for Sanger sequencing and of siRNA targeting EIF4H.

**Table S7** Ribo-seq and RNA-seq coverage -related to Figure S3-.

